# oncotree2vec – A method for embedding and clustering of tumor phylogenetic trees

**DOI:** 10.1101/2023.11.16.567363

**Authors:** Monica-Andreea Baciu-Drăgan, Niko Beerenwinkel

**Author notes:** **Code availability:** Oncotree2vec is implemented in Python and freely available at https://github.com/cbg-ethz/oncotree2vec.

## Abstract

Understanding the genomic heterogeneity of tumors is an important task in computational oncology, especially in the context of finding personalized treatments based on the genetic profile of each patient’s tumor. Tumor clustering that takes into account the temporal order of genetic events, as represented by tumor mutation trees, is a powerful approach for grouping together patients with genetically and evolutionarily similar tumors and can provide insights into discovering tumor sub-types, for more accurate clinical diagnosis and prognosis. Here, we propose oncotree2vec, a method for clustering tumor mutation trees by learning vector representations of mutation trees that capture the different relationships between subclones in an unsupervised manner. Learning low-dimensional tree embeddings facilitates the visualization of relations between trees in large cohorts and can be used for downstream analyses, such as deep learning approaches for single-cell multi-omics data integration. We assessed the performance and the usefulness of our method in three simulation studies, and on two real datasets: a cohort of 43 trees from six cancer types with different branching patterns corresponding to different modes of spatial tumor evolution and a cohort of 123 AML mutation trees.

## 1 Introduction

Understanding the genomic heterogeneity within and among tumors is key for personalized oncology approaches. A therapy that works for one patient might also be effective for another patient with a similar tumor profile. Therefore, tumor clustering based on the genetic patterns of tumors can provide insights into discovering tumor subtypes for a more accurate clinical diagnosis and prognosis.

Tumors arise from single cells that undergo diverse genomic aberrations, leading to diverging malignant clonal lineages which result in distinct subpopulations. Recent developments in single-cell omics technologies, which can measure different tumor modalities (genomics, transcriptomics, etc.) at the single-cell level, have revealed, with increasing precision, intra- and inter-tumor heterogeneity, including the evolutionary history of each tumor and its cell phylogeny.

Several methods to discover cancer subtypes and tumor clusters have been proposed, based on clustering data modalities individually [2, 8, 12], or using multi-omics integrative approaches [51, 1, 5]. However, clustering tumors based on their evolutionary histories has, to our knowledge, not been explored before.

The evolutionary history of a tumor can be represented by a mutation tree. It captures the evolutionary relationships among different tumor subclonal populations at the time of the biopsy, i.e., the (partial) temporal order in which the genomic aberrations were acquired [4], which can be used to predict tumor progression and clinical outcome [32, 41, 42]. At the same time, the different shapes and branching patterns of the mutation trees reflect different modes of tumor evolution governed by selection and by the spatial architecture of the tumor [39].

Tumor mutation trees are rooted trees where the nodes represent genomic events such as point mutations, copy number alterations (CNA), or other sets of genomic alterations, and the nodes are connected according to their order in the evolutionary history. Mutation trees can be obtained from single-cell data using several tools, such as SCITE [23], OncoNEM [43] and and HUNTRESS [28] for point mutation trees, SCICoNE [27] and CONET [33] for copy number trees, or COMPASS for joint copy number and point mutation trees [47].

Existing methods for finding similarities between mutation trees are based on graph metrics that consider single handcrafted features or individual specific patterns in the graphs, such as the number of edge-induced partitions [6, 22], conserved regions or evolutionary trajectories [21, 7], matching node triplets [9], common ancestor or distinctly inherited sets [14], or weighted Pearson’s correlation scores for the mutational characteristics of each subclone [44].

However, all these metrics are based on the infinite sites assumption (i.e., mutations occur only once and never disappear), which is questionable for point mutation trees [25] and definitely inappropriate for copy number trees, where amplifications and deletions affecting the same gene occur frequently. Moreover, each of these distance metrics looks for specific tree patterns, and there is no existing method to combine these different basic metrics in order to discover more complex patterns that characterize real tree cohorts.

More complex approaches for graph similarity measures and clustering include learning vector representations of graph nodes or whole graphs using graph kernels [36] or Graph Neural Networks [20]. However, none of the existing methods take into consideration the biologically relevant node relations inside the tumor mutation trees such as co-occurring mutation pairs (i.e., mutations in a common lineage), or clonally exclusive mutations (which appear in different branches) [3, 13, 19, 26].

Here we propose oncotree2vec, to our knowledge the first unsupervised learning method for learning vector representations (*aka* embeddings) that capture the different relationships among subclones, with the purpose of mutation tree clustering. Our method matches trees based on both tree node neighborhoods of different degrees and biologically meaningful pairwise relations between the tree nodes that correspond to clonal co-occurrence and exclusivity. We visualize the clustering results by highlighting the matching subtree structures which dominate each cluster.

We apply our method to three datasets generated in simulation studies and two real datasets: one cohort of 43 trees of different branching patterns from six cancer types corresponding to different modes of tumor evolution [39], and another cohort of 123 AML mutation trees [35].

## 2 Methods

We developed oncotree2vec, a method for embedding and clustering mutation trees (Fig. 1). The nodes are labeled by sets of point mutations, copy number alterations, or any finite subset of genomic events. We use a Neural Network (NN) language model to learn, in an unsupervised manner, vector representations of mutation trees that uniquely capture the different relationships between subclones across the entire cohort so that the trees with matching subtree structures and labels have a close vector representation in the embedding space. The learned embeddings produced by oncotree2vec can be further used for clustering and similarity heatmap visualization of the relations between the mutation trees in the cohort using metrics such as, for example, the cosine distance. The results of the clustering are interpretable by inspecting the matching tree labels and sub-structures which dominate each cluster.

**Figure 1.**
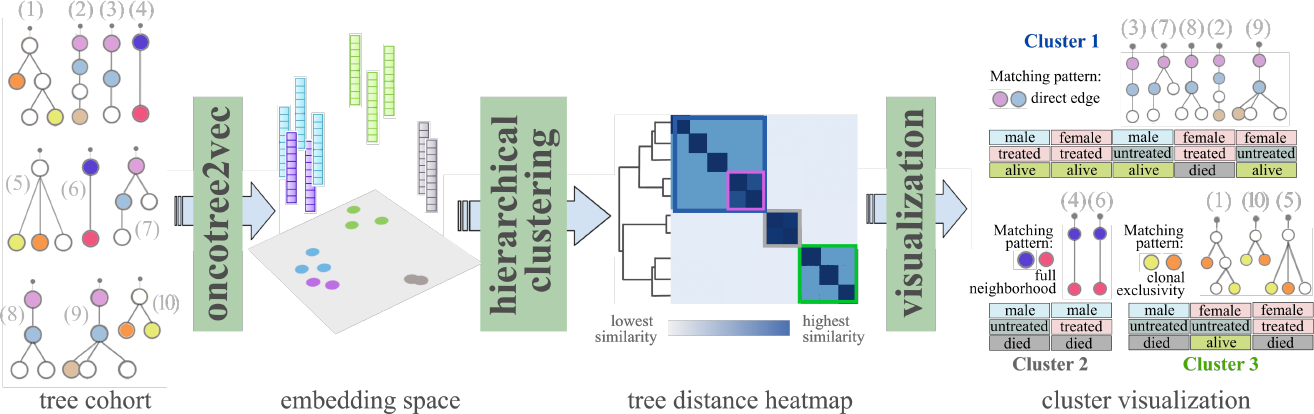
Embedding and clustering of mutation trees using oncotree2vec. Oncotree2vec learns vector representations of every tree in a cohort such that the trees with matching patterns will have similar vector representations in the embedding space. The embeddings are then used for clustering and heatmap visualization. The colors of the vectors in the embedding space correspond to the colors of the squares indicating the clusters in the tree distance heatmap. The results can be interpreted by inspecting the matching subtree structures which dominate each cluster through a javascript visualization. The tree node colors indicate matching node labels. In this example, the first cluster contains trees 3, 7, 8, 2 and 9, which share a direct edge, and correspond to the blue square on the heatmap. Furthermore, trees 2 and 9 share an additional co-occurring leaf node and form a sub-cluster (the violet square on the heatmap). Cluster 2 contains the identical match of trees 4 and 6 (the gray square on the heatmap). Finally, the third cluster contains trees 1, 10 and 5, which share a pair of clonally exclusive nodes.

### 2.1 Tree embedding learning using a neural document embedding model

Oncotree2vec is inspired by graph2vec [36], an approach for learning labeled graph embeddings in an analogous fashion to doc2vec [29], a NN language model for document embeddings. Doc2vec is based on the idea that words which appear in similar contexts tend to have similar meanings and hence should have similar vector representations. Given a set of documents *D* = {*d*_1_, *d*_2_, …, *d*_*N*_ } and sequences of words *c*(*d*_*i*_) = {*w*_1_, *w*_2_, …, *w*_*Li*_} sampled from each document *d*_*i*_ ∈ *D*, where *L*_*i*_ is the number of words contained by *d*_*i*_, the model learns a *δ*-dimensional embedding *emb* of the document *d*_*i*_ and of each word sampled from *c*(*d*_*i*_), i.e., *emb*(*d*_*i*_), *emb*(*w*_*j*_) ∈ *R*^*δ*^. Once the training algorithm converges, the learned vector representations are the weights of the last hidden layer of the NN which correspond to the input words and to the input document. The optimization function of the NN maximizes the log likelihood,

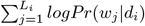

where *Pr*(*w*_*j*_| *d*_*i*_), the probability that target word *w*_*j*_ appears in the context of the document *d*_*i*_, is defined as

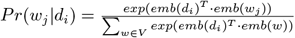

and V is the vocabulary of all words across all documents in D. The matching words and documents are mapped to a similar position in the embedding space, while dissimilar ones are pushed away from each other.

In the case of mutation trees, each tree is uniquely described by the set of labeled subtree structures, referred to as the *vocabulary*. Like graph2vec, oncotree2vec considers the tumor mutation trees analogously to documents and the labeled subtree structures analogously to words from a specific vocabulary, and adapts the document embedding model to learn embeddings for tumor mutation trees.

Similar to graph2vec, oncotree2vec extracts for each mutation tree a set of labeled rooted subtrees of different neighborhood sizes (starting with size 0) around each node, which are encoded as vocabulary words using the Weisfeiler-Lehman (WL) kernel [49]. The WL subtree kernel is based on iteratively decomposing the trees into rooted subtrees around every node and relabeling these nonlinear structures by aggregating the labels of a node and its direct neighbors into a string that is hash compressed (Suppl. Fig. 1). Applying this procedure iteratively generates labels encoding increasingly larger neighborhoods of each node, allowing to compare more extended nonlinear tree substructures. In addition, oncotree2vec takes into consideration the biologically relevant subclone relations in tumor mutation trees and expands the tree vocabulary by adding words corresponding to pairs of nodes of interest, specifically, pairwise relations that capture direct or indirect clonal lineages, mutually exclusive pairs, root-child relations and words that encode the subtree structures, where the labels are discarded (Fig. 2). For encoding the subtree structures we apply the same WL kernel strategy, but on the unlabeled trees, and discard the individual nodes (kernel size greater or equal to 1).

**Figure 2.**
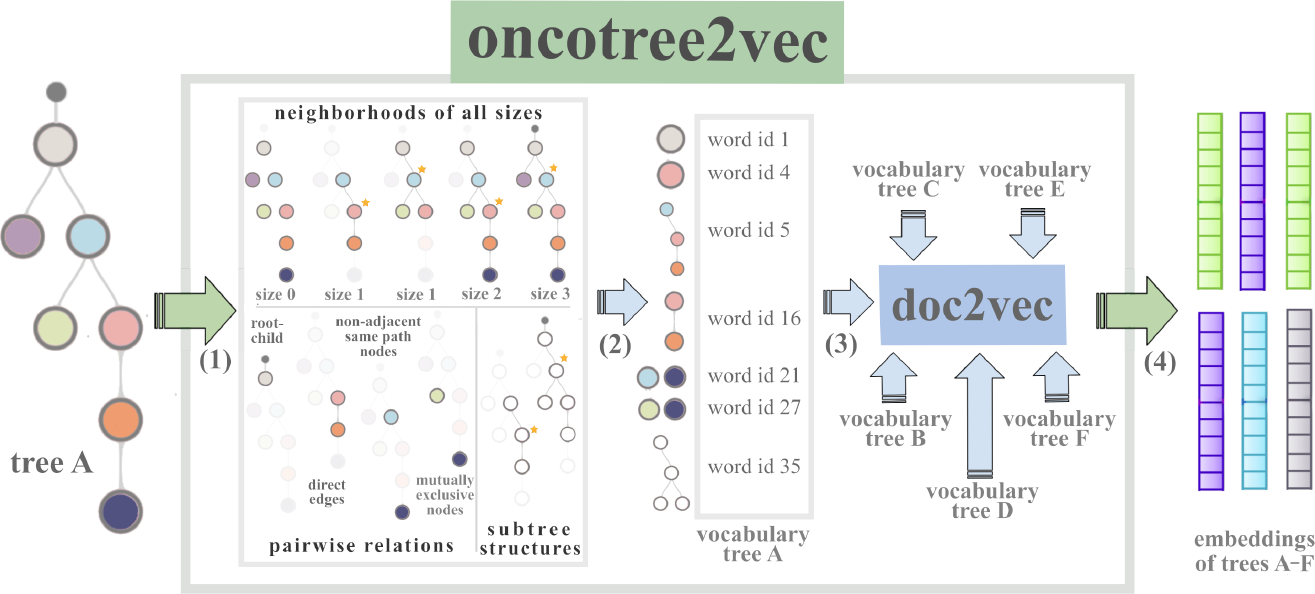
Oncotree2vec internal workflow. Each tree is described by the set of its subtree structures that captures the different relationships between the subclones of the tree, namely, labeled subtrees of different neighborhood degrees around each node, pairwise relations that capture direct or indirect clonal lineages, mutually exclusive pairs, and subtree structures, where the labels are discarded. The neighborhoods of size 0 are the individual nodes. Selected examples of subtree structures are shown for **step (1)**. The subtree structures are encoded by numerical ids and used as vocabulary words – **step (2)**. Each input tree, encoded by its vocabulary, is input to doc2vec, which uses a NN language model to produce the vector representations corresponding to each of the original trees – **step (3)**. The trees with matching patterns will have a close vector representation in the embedding space – **step (4)**.

The embedding learning process is based on the idea that if a word appears in the vocabulary of a pair of trees, their corresponding embeddings become more similar in the embeddings space. Similarly, for every non-matching word in the vocabulary of two trees, the corresponding embeddings are pushed away from each other. Therefore, by reducing the number of unique words in the vocabulary, we increase the chance of weak similarities to be detected. This is why for pairwise relations, which have high biological relevance, we only include the words which are non-unique across the entire cohort. That is, if a tree contains a certain pair of labeled nodes which is unique at cohort level, we do not add the corresponding word to the vocabulary of that tree.

We trained our model using doc2vec with negative sampling, employing an adapted tensorflow implementation from graph2vec [36]. During training we add duplicates of every tree (and therefore double the number of trees) in order to enforce the network to learn the identical match. The expected distance between the learned embeddings of duplicate trees is 0. The learned distance between duplicate trees is used as an indicator for choosing a cutoff number of training iterations.

### 2.2 Model parameters

Oncotree2vec is a versatile framework, allowing the user to adjust certain parameters of the model. The model has two sets of adjustable parameters, which come into play at different steps of the algorithm. First, each category of node relations (individual nodes, neighborhoods around the nodes, root-child relations, direct edges, non-adjacent same-path pairwise relations, mutually exclusive pairs and tree structures; see Fig. 2) can be weighted in order to guide the machine learning algorithm on which patterns to include or to augment in the vector representations of the input mutation trees. Second, some machine learning-specific parameters need to be tuned in order to reach an optimal solution.

Our model extracts subtree structures to build a vocabulary for every tree in the cohort. However, by default some of the vocabulary categories are overrepresented, and others are underrepresented. On the one hand, encoding neighborhoods of different sizes around every node using the Weisfeiler-Lehman (WL) kernel generates a large amount of words, namely the number of tree nodes times the size of the largest WL kernel applied, per tree (Suppl. Fig. 1). On the other hand, for words corresponding to the vocabulary categories corresponding to pairwise relations (i.e., root-child relations, direct edges, and node pairs corresponding to clonal co-occurrence and exclusivity), we only include words which are non-unique across the entire cohort. Therefore, the number words encoding the pairwise relations are underrepresented compared to the number of words corresponding to node neighborhoods, a situation that can be adjusted by augmenting the number of underrepresented words. For example, for an augmentation of 2x of the root-child relations, we double the number of words corresponding to the root-child relations in the vocabulary of every tree. As a result, the root-child relations would weight twice as much in the tree matching process. Similarly, depending on the purpose of the tree clustering, every tree vocabulary category can be ignored or weighted by different amounts when constructing the vocabulary, using the augmentation parameter of the model (see the different augmentation amounts from Suppl. Table 1).

The WL kernel size can also be specified by the user. In practice, it is very unlikely to find large complex labeled subtrees that fully match in real data. In general, a maximum WL kernel size of 3 is enough to distinguish unique matches (see the examples in Suppl. Fig. 1 and Suppl. Fig. 2A, where the WL kernel sizes of 2 and 3 respectively cover the whole tree).

Another set of parameters that need to be set in order to reach an optimal solution are the machine learning-specific parameters, namely, the embedding size and the number of training iterations. By default we use a standard embedding dimension of *δ* = 128. However, for small cohorts, e.g., smaller than 100 samples, using a large embedding size results in sparse distribution of the vector representations in the embedding space, which gives weaker similarity between the learned embeddings and weaker separation between the clusters (Suppl. Fig. 5C). On the other hand, an embedding size which is too small is not able to encode all the tree patterns, giving a weak separation between the tree embedding clusters (Suppl. Fig. 5A). Therefore, for small cohorts it is recommended to run the model with a lower embedding size, e.g., *δ* = 64 or 32.

During training we assess various quantities to decide when to stop the training and select the optimal solution to avoid overfitting. First, we plot the residual at every training iteration in order to determine when the curve of the convergence plot becomes steady, i.e., the model converges. We also measure after every iteration the following parameters that show how the learned embeddings span over the entire embedding space: the minimum and maximum cosine distance between the learned embeddings (a fully covered embedding space would contain pairs of embedding vectors for which the cosine distance covers the whole range between 0 and 1) and the maximum cosine distance between duplicated trees (which is expected to be 0).

## 3 Results

To demonstrate the effectiveness of oncotree2vec, we assess its performance on three synthetic datasets of simulated trees designed to answer the following questions: (1) Does the clustering using the learned embeddings correspond to the expected tree cluster separation? (2) Do the pairwise distances between the learned embeddings correlate with the known tree rank ordering constructed by design? (3) How do the similarities learned using the oncotree2vec embeddings compare to standard tree distance metrics?

In addition, we assess the performance and the usefulness of our method on two tasks involving real data: first, we apply oncotree2vec to cluster tree structures and distinguish different modes of tumor evolution; and second, we discover subgroups of patients with matching genomic evolutionary patterns, potentially leading to a better stratification of tumors with respect to survival. We apply oncotree2vec to a cohort of trees with different branching patterns from six cancer types, corresponding to different tumor spatial evolutions [39] and to a cohort of 123 AML mutation trees inferred using SCITE [23] from single-cell DNA sequencing data [35]. An overview of all mutation tree cohorts we analyzed is presented in Table 1. We show the model parameters used in each of the experiments in Suppl. Table 1. The reported results correspond to a cutoff number of iterations after the training reaches convergence (i.e., the curve of the convergence plot becomes steady).

**Table 1:** Overview of the mutation tree cohorts used in the experiments. For the trees from the third experiment (real trees with different modes of evolution) we are interested only in the tree structures, therefore reporting the number of mutations is not applicable.

| Experiment | Number of samples | Total number of clones | Average number of nodes per sample | Average node degree | Number of mutations |
| --- | --- | --- | --- | --- | --- |
| Synthetic dataset I | 315 | 2,331 | 7.4 | 1.72 | 108 |
| Synthetic dataset II | 52 | 2,600 | 50 | 1.96 | 1,325 |
| Synthetic dataset III | 26 | 208 | 8 | 1.62 | 26 |
| Real trees with different modes of evolution | 43 | 667 | 15.5 | 1.87 | NA |
| AML mutation trees | 123 | 641 | 5.2 | 1.8 | 31 |

### 3.1 Simulation studies

#### 3.1.1 Clustering a cohort of 16 groups of synthetic trees

For this experiment we simulated trees with known clustering and known order of the pairwise similarities between the trees, referred to as rank ordering. First, we simulated 16 groups of trees with different branching, sizes, and labelings (Fig. 3A), so that the trees within the same group are more similar to each other in terms of tree structure and labeling than any other tree from the other groups. Each group contains trees with identical structure and node labeling variations starting from a tree with an initial labeling. The labels of the subsequent trees from the same group match the corresponding labels of different neighborhoods with decreasing sizes from the start tree (see an example in Suppl. Fig. 2A). Therefore, the larger the start tree, the larger the group (the group size ranges from 9 to 34 trees; Table 1, synthetic data I). The rank ordering of the trees inside each group is induced by the number of elements of the intersections between the vocabularies of each pair of trees in the group (i.e., the number of sub-tree structures and labels that each pair of trees has in common). The fixed tree structures per group ensure the same size of vocabulary for trees inside the same group and allow us to focus on the similarity rank ordering of the trees induced by the labelings.

**Figure 3.**
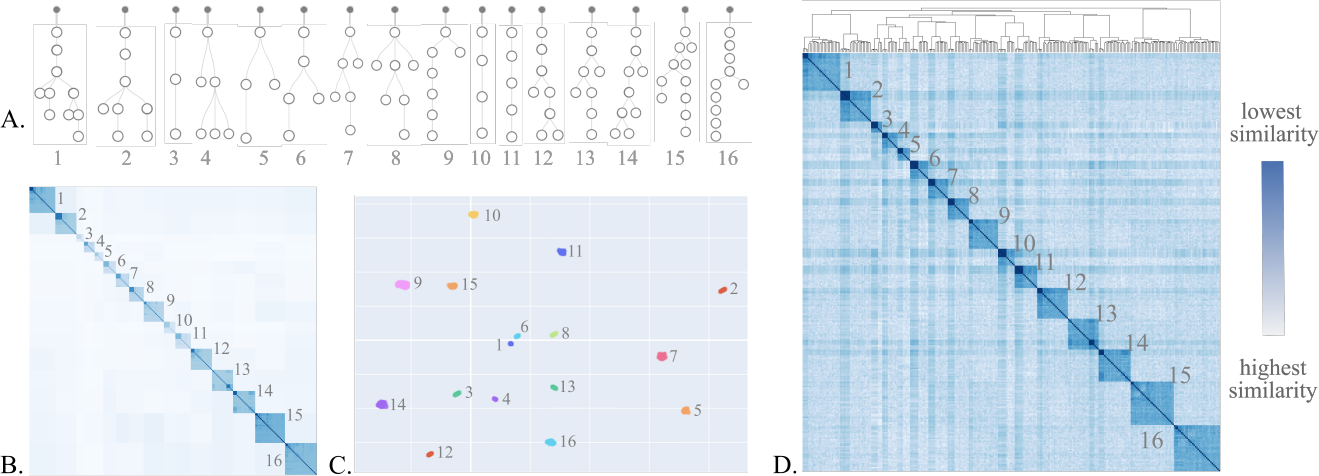
Clustering a cohort of 16 groups of synthetic trees (synthetic dataset I). **(A)** Tree structures of the start trees for each of the 16 groups. **(B)** Heatmap showing the number of elements of the vocabulary intersections for each pair of trees. White color corresponds to empty intersections, and dark blue indicates large vocabulary intersections. **(C)** UMAP showing the separation between the inferred tree clusters using the cosine distance between the learned embeddings. **(D)** Heatmap visualization of tree similarities for the simulated trees from 16 groups, using the cosine distance between the embeddings learned by oncotree2vec. The trees cluster by simulation group. (B-D) The numbers correspond to the tree indices in panel A and indicate the tree structure of the trees in each group.

We train oncotree2vec on the simulated trees from the 16 groups, each tree being represented by a vocabulary that takes into consideration the tree structures and the labels of the neighborhoods of different sizes, and cluster the learned embeddings using the hierarchical clustering with cosine distance metric (a standard approach for measuring embedding similarities in the machine learning field) and ward linkage method. We evaluate the output after 7,000 iterations, when the training starts to converge, as shown in Suppl. Fig. 3A, B. The results show that the trees from the same group cluster together and that the distance between the learned embeddings mimics the size of the vocabulary intersections for each pair of trees (Fig. 3B, D). The clusters are segregated by group with a silhouette score of 0.384 (Fig. 3C, Suppl. Fig. 3A).

The weaker similarities between different clusters is explained by the fact that, even if the tree structures are designed to be different from one group to the other, there are still small subtree structures which are matching across different groups. However, these similarities are very few compared to the large number of similar patterns between the trees from the same group. We obtained an average silhouette score of 0.374 after 10 runs of the same experiment with optimal number of iterations.

Next, we compute the deviation of the embedding similarities from the given order inside each cluster. The similarity scores of each tree w.r.t. the start tree are ordered by the number of vocabulary intersections in decreasing order (Suppl. Fig. 2B, C). Each similarity score should be lower or equal to the minimum score from the previous rank (see an example in Suppl. Fig. 2D). The deviation of the embedding similarities from the expected order is computed by subtracting, for each tree, the computed similarity w.r.t. the start tree from the minimum similarity score from the previous rank. The optimal deviation is 0. We compute normalized deviation from the expected ranking of the embedding similarities inside each group of simulated trees after different numbers of iterations and find a normalized deviation across all the 16 clusters of 0.0089 after the training reaches convergence (cutoff at 7,000 iterations)– Suppl. Fig. 3B,C.

#### 3.1.2 Clustering synthetic trees with known similarity rank ordering

For the second experiment of generating synthetic data, we construct a cohort of 52 linear trees of 50 nodes each (one start tree, 50 trees with matching nodes w.r.t. the start tree, and one tree without matching nodes; Table 1, synthetic dataset II) labeled such that the intersection between the corresponding tree vocabularies ranges between 0 and 50. In order to achieve this, for simplicity, we start from a reference linear tree with 50 nodes, each node having a different label, and derive trees with the same structure and decreasing number of matching nodes with respect to the initial tree (Suppl. Fig. 4A). In order to obtain the desired number of matches between the pairs of trees we build the vocabulary based on the labels of the individual nodes and discard all the other tree patterns.

Again, we expect that the pairwise distances between the embeddings learned by oncotree2vec (Suppl. Fig. 4C) reflect the known tree rank ordering based on the number of shared matches between the pairs of trees, which was set when constructing the synthetic trees (Suppl. Fig. 4B). As before, we compute the mean deviation of the embedding similarities from the expected ranking with respect to each of the 50 trees, i.e., we take each of the trees as a reference tree and compute the cosine distances between the embedding of the reference tree and all the other trees (Suppl. Fig. 4D). The lowest average deviation value of 0.02 corresponds to the embeddings learned after 200 iterations and corresponds to the moment when the training starts to converge (Suppl. Fig. 4F).

In addition, we also evaluate the consistency of the learned embeddings for the tree pairs with the same number of matching nodes, which should be similar – they appear on the antidiagonals parallel to the main antidiagonal of the heatmap in Suppl. Fig. 4B. In this respect, we expect the pairwise cosine distances between the learned tree embeddings corresponding to pairs of trees with the same number of matching nodes to be very close to zero. We find a standard deviation value of 0.02 after the training reaches convergence (cutoff at 200 iterations), which is close to zero, as expected (Suppl. Fig. 4E, F).

#### 3.1.3 Comparing oncotree2vec to graph2vec and to other tree distance metrics

The tree embeddings learned by oncotree2vec can be used to match biologically relevant patterns in tree cohorts. We can obtain a distance metric by applying the standard cosine distance on the tree embeddings learned by applying oncotree2vec on a cohort of trees and use this score to compare our results to other methods for computing pairwise tree similarities. We build a dataset of 13 selected pairs of trees that are compared two by two, each pair containing one precise matching pattern from each vocabulary category (Table 1, synthetic dataset III). Ideally, all pairs of trees sharing one pairwise relation should receive the same distance score. One tree pair with a matching neighborhood of size 1 should get the strongest match out of all the tree pairs (it contains multiple combined matching patterns: one neighborhood of size 1 and three co-occurring mutation pairs). Finally, we also add two pairs of trees which do not match because of the reverse order of the mutation events, and the different branching of the events, respectively (the corresponding distance score should be 0).

In Suppl. Table 2 we show the pairwise distance scores for the selected pairs of trees computed using oncotre2vec and compare them to CASet and DISC [14], MP3 [9], Bourque distance [23] and graph2vec [36]. We found that all the metrics except CASet and Bourque find the strongest match (i.e., the minimum distance) for the pair of trees matching one neighborhood of size one (first tree in Suppl. Table. 2). Furthermore, we find that the CASet and DISC metrics are sensitive to the location of the matching nodes w.r.t the root. On the other hand, MP3 meets our expectation and reports equal distances for all the pairs of trees matching pairwise relations, but it fails to account for root-child relations and has a frequency distribution of distances biased towards higher distance scores, which was also reported in [23]. Similarly, the tree distance scores obtained using the embeddings learned by graph2vec in two different settings (WL kernel size of 1 and 2) do not highlight the matching patterns between the pairs of trees, giving close scores between the pairs expected to match and all the non-matching trees in the tree set (silhouette scores 0.14 and 0 respectively). Both the result of the triplet-based approach (MP3) and the one obtained with graph2vec have a similar explanation: the larger the tree, the higher the number of all possible triplets and node neighborhoods. Therefore the only matching pattern underlying the pairs of trees is underrepresented in the set of all patterns that count for the matching. This problem is solved with oncotree2vec by the customized weighing of each vocabulary category and by removing from the tree vocabularies the unique patterns of pairwise relations among the entire cohort. The results given by oncotree2vec are the closest to our expectation, obtaining a good separation between the matching groups with and without including the node neighborhoods in the vocabulary, reflected in silhouette scores of 0.4 and 0.839, respectively.

### 3.2 Highlighting different models of tumor evolution on real data

Next, we apply oncotree2vec to real data for identifying different tumor growth patterns corresponding to different modes of spatial tumor evolution. We use a cohort of 43 tumor mutation trees collected in [39] from different publications, for six cancer types: acute myeloid leukemia (AML, [35] – 8 samples), clear cell renal cell carcinoma (ccRCC, [50] – 5 samples), mesothelioma ([55] – 6 samples), breast cancer ([34] and [54] – 11 samples), non-small cell lung cancer ([24] – 5 samples), and uveal melanoma ([15] – 8 samples).

By using oncotree2vec to cluster the corresponding tree structures we collect the optimal output after 500 iterations (Fig. 4B) and distinguish four clusters (Fig. 4A) which match the four established modes of tumor evolution as follows. The first cluster contains trees with a small number of nodes and mostly linear tree shapes (cluster 1 in Fig. 4C). It comprises samples from two cancer types: AML and mesothelioma, both corresponding to linear evolution because of their unconstrained spatial evolution (encountered in liquid tumor evolution) and slow cell turnover due to few driver mutations (specific to mesothelioma), as shown in [12, 40].

**Figure 4.**
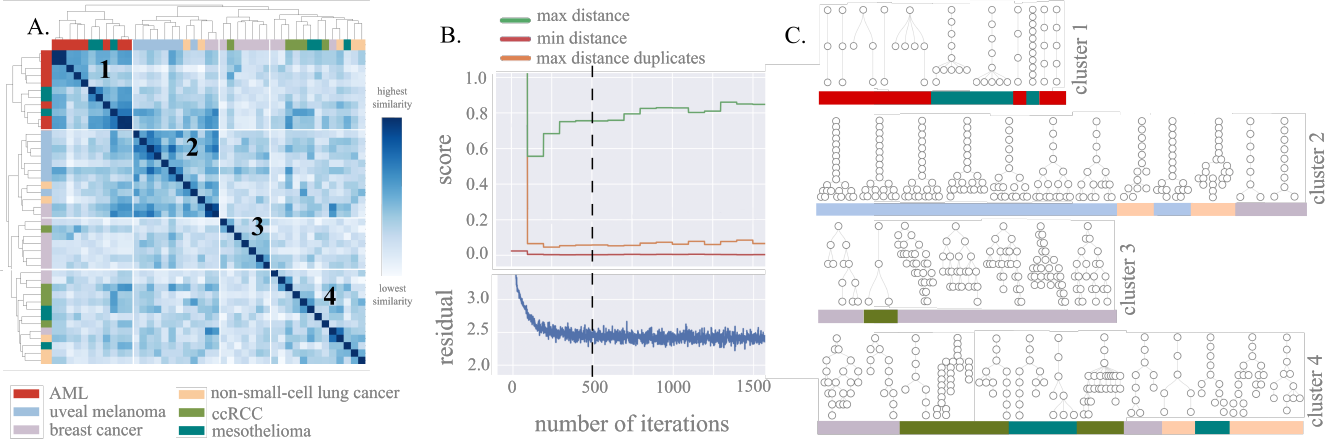
Clustering mutation tree structures from six cancer types. **(A)** Heatmap of tree similarities for 43 tumor mutation trees from 6 cancer types, using the cosine distance between the embeddings learned by oncotree2vec. **(B)** Training parameters: minimum and maximum cosine distance between the learned embeddings, the maximum cosine distance between duplicated trees (see Section 2.2) and the residual after each training iteration. The dotted line shows where the convergence plot becomes steady, i.e., the training algorithm starts to converge (this is the cutoff used). **(C)** Visualization of the tree structures of the tumor mutation trees in each of the four identified clusters corresponding to four modes of evolution: linear, punctuated, branching and linear-to-branched evolution. The length of lines connecting the tumor subclones and the node sizes are not informative.

The second cluster displays structure patterns specific to punctuated tumor evolution (cluster 2 in Fig. 4C), where the tumor starts with a linear evolution and then a large number of genomic aberrations occur in a short amount of time [11]. This is also confirmed by the cancer type dominating this cluster, namely uveal melanoma, which has been described to follow this mode of evolution [18]. The third cluster contains trees that follow branching evolution (cluster 3 in Fig. 4C), which produces trees in which clones diverge and evolve in parallel. Such modes of evolution have been reported in studies of breast cancer [38, 46], which indeed dominates this cluster.

The remaining trees from the cohort cluster together in a fourth cluster of tumors which exhibits patterns of both linear and branching evolution (cluster 4 in Fig. 4C). A possible biological interpretation of this behavior is related to the fact that the evolution of real-world tumors may be influenced by both global and local restrictions. Global restrictions (such as tumor size) increase the overall selection pressure and cause an increase in the number of driver mutations, while local growth restrictions (caused by local tissue structures such as glands) increase clonal diversity. A combination of the two factors can result in mixed evolutionary patterns of both linear and branching evolution, known as linear-to-branched evolution [48]. These transitions between evolutionary modes were observed in breast and lung cancers, which exhibit localized growth into separate mammary glands and alveoli, respectively [48, 30], but also in many colorectal cancers [39, 52], which contribute to the fourth cluster.

### 3.3 Clustering a cohort of 123 AML mutation trees

Finally, we apply oncotree2vec to the entire cohort of 123 AML mutation trees inferred from single-cell DNA sequencing data from [35], and analyze different tumor evolution patterns present in the dataset (for longitudinal samples, we only consider the latest biopsy of each patient). First, we assess the tree structures without taking into consideration the subclonal mutations and confirm both the linear and branching clonal evolution patterns present in AML, which were discussed in [35] (Fig. 5A,C).

**Figure 5.**
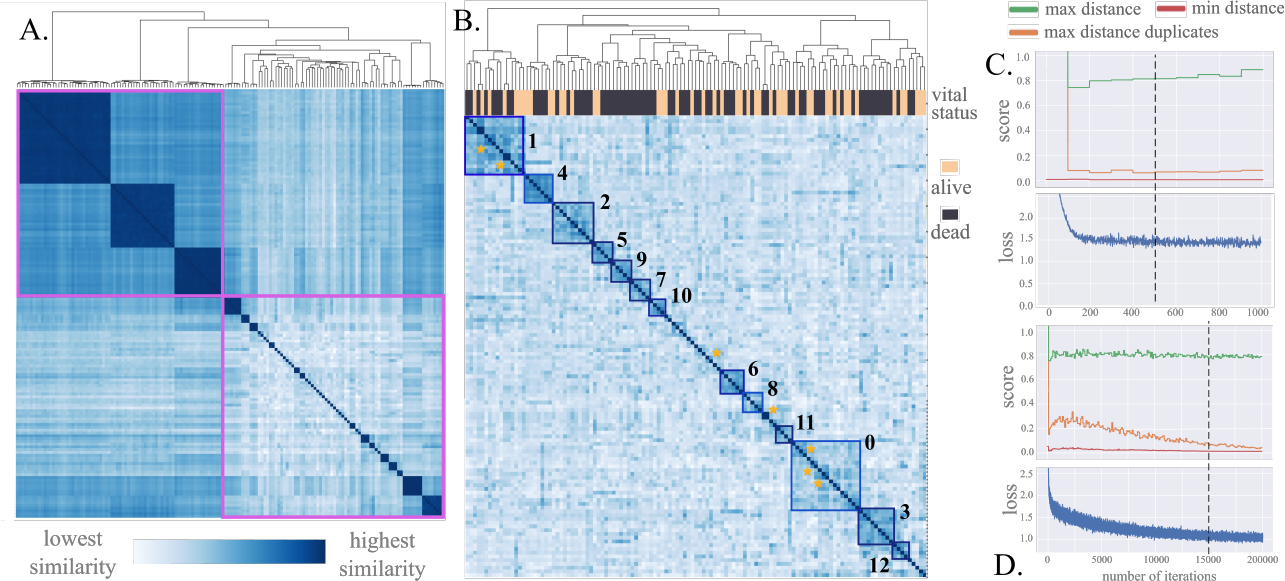
Figure 7. Clustering results for a cohort of 123 AML mutation trees. **(A)** Heatmap visualization of tree similarities based on the tree structures. The cluster split indicates two different modes of clonal evolution: linear and branching. **(B)** Heatmap visualization of tree similarities based on the individual sub-clone labels and different labeling relations: neighborhoods of different sizes, rootchild relations, direct edges, non-adjacent same path pairwise relations and mutually exclusive relations. The large clusters are indicated by squares, and small clusters are highlighted by a star. The red squares indicate clusters of patients with high average survival or death rate. **(C, D)** Training parameters for the two clusterings: minimum and maximum cosine distance between the learned embeddings, the maximum cosine distance between duplicated trees (see Methods 2.2) and the residual after each training iteration. The dotted line shows where the convergence plot becomes steady and the training algorithm starts to converge (this is the cutoff used).

Next, we train oncotree2vec embeddings that encode information about the individual subclone labels and different labeling relations: neighborhoods of size up to 3, root-child relations, direct edges, clonal co-occurrence and clonal exclusivity, and identified AML patient groups with matching patterns. Specifically, after the convergence of the model (at 15,000 iterations – Fig. 5D) we found 13 clusters of sizes between 18 and 4 samples governed by a primary gene mutation, captured by the model through the root-child relations (the squares in Fig. 5B) and 7 strong relations of clonal co-occurrence and exclusivity among groups of 4, 3, or 2 samples (starred in Fig. 5B). Details about each of the clusters can be found in Suppl. Table 3. We also performed survival analysis to identify high/low risk groups (Suppl. Fig. 7).

For AML, the primary gene mutation is a strong signal, known to be the main factor to drive the development of AML, and it is used as the first criterion for diagnosis in clinical protocols [16]. By employing survival analysis we found two groups of clusters with significant difference in survival (p-value *<* 0.05 in the pairwise log-rank test): cluster 1, governed by the primary mutation IDH2, vs the group of clusters 3, 4, 6, and 9 with primary mutations NRAS, TP53, SF3B1, FLT3-IDT. The difference in survival is confirmed by the separation of Kaplan Meier curves (Suppl. Fig. 7) and by the median survival time of 55.3 months for cluster 1 and 9-14 months for the group of 4 clusters (Suppl. Table 3). The better prognosis in cluster 1 might be confirmed by the fact that IDH2, the primary mutated gene for the patients in cluster 1, is not part of a distinct prognostic group according to the official clinical recommendations for diagnosis in 2022 [16].

For smaller clusters of 2 or 3 patients, we also identified further matching patterns of co-occurrence and clonal exclusivity among functionally redundant mutations, namely IDH1 and IDH2, among genes involved in receptor tyrosine kinase (RTK)/Ras GTPase (RAS)/MAP Kinase (MAPK) signaling pathway (FLT3, NRAS, KRAS, PTPN11), and other local patterns described in Suppl. Table 3 which correspond to previously published reports [35, 31].

## 4 Discussion

We have developed oncotree2vec, an unsupervised representation learning technique to learn vector representations of tumor mutation trees that uniquely capture the different relationships between tree subclones by taking into consideration biologically meaningful pairwise relations of clonal co-occurrence and exclusivity. Our method can be used for visualization and clustering of mutation tree cohorts, which has, to our knowledge, not been addressed before.

We evaluated the performance of our method on two synthetic datasets of trees with known clustering and similarity rank ordering based on the number of matches shared between the trees, and showed that trees with matching subtree structures and labels have a close vector representation, while dissimilar ones are pushed apart in the embedding space. We also evaluated how the similarities learned using the oncotree2vec embeddings compare to standard tree distance metrics using a third synthetic dataset. We found that the results obtained with oncotree2vec were the closest to our expectation and highlighted the weaknesses of the other methods.

Next, we demonstrated the usefulness of oncotree2vec on two real-world cohorts of mutation trees. Using the embeddings learned by oncotree2vec we first clustered the mutation trees from a cohort of six cancer types based on patterns in their tree structure and identified four different modes of evolution: linear, punctuated, branching and linear-to-branched evolution that are in line with existing literature for each cancer type. Then we clustered a cohort of 123 point mutation trees based on their matching evolutionary patterns and found 13 clusters governed by a primary gene mutation and 6 strong relations of clonal co-occurrence and exclusivity among smaller groups of 4 samples or less. We were able to run all the experiments on a desktop PC in less than 3 hours.

Both our experiments and existing literature [35, 50, 24] shows that tumor mutation trees from real data cohorts typically do not share complex patterns, potentially identifiable with larger WL kernel sizes. This is because of the known high levels of heterogeneity in cancer tumors caused by the tumor spatial architecture and the resistance to treatment [10]. For example, no large clusters of trees with large neighborhoods matching were found in the AML dataset, thus the need of matching additional patterns describing pairwise relationships between clones, which oncotree2vec addresses (Suppl. Fig. 6).

The performance of oncotree2vec also depends on the way the mutation trees are constructed. We use the order of the acquired mutation events to learn tree embeddings that match biologically meaningful pairwise relations of mutation trees. Therefore, the performance of oncotree2vec is highly influenced by the resolution at which the trees can be inferred (which depends on the number of single cells analyzed, amplification biases and errors) and by the accuracy of the tree reconstruction method, and the relevance of the results depends on that.

Our experiments show the consistency of the embeddings for the pairs of trees with the same number of matching patterns inside the same cohort and the stability of the embedding among multiple runs. Also, our simulation studies show that the pairwise similarities between trees follows the rank ordering induced by construction of the synthetic dataset. However, the main challenges of applying unsupervised deep learning methods in genomics are the limited amount of training data, the high number of parameters to tune and the risk of overfitting and the lack of interpretability [53, 45]. However, in Section 2.2 we provided solutions for tracking and improving the performance of our training algorithm. We added the capability of tracking the steadiness of the residual function and the minimum and maximum cosine distance scores between control samples in the cohort for choosing the optimal number of iterations, we provided the possibility of augmenting underrepresented words in the tree vocabulary in order to highlight the weak matches inside the cohort and suggested reducing the embedding size for smaller cohorts. Also, despite being an unsupervised learning method, oncotree2vec provides interpretable results due to the fact that the tree embedding similarities reflect the intersection of vocabulary subtree structures and each cluster of mutation trees can be summaries by the set of matching subtree structures which dominate the the cluster, therefore the lack of interpretability that most of the unsupervised deep learning have is not an issue for oncotree2vec.

Also, depending on the input dataset, there is no guarantee that the data admits a perfect clustering, either because the data may not come from well-defined clusters, or because the clustering algorithm is not able to efficiently find well separated stable clusters, which can be the case when the metric used for computing the similarities between embeddings is not transitive (as is the case, for example, for the cosine distance), or when the optimization function does not converge to a stable result. However, the key limitation of these methods is that trees cannot be reliably embedded into inner product space or an Euclidean space of low dimensions, and more complex hierarchical latent structures might be more suitable for representing the data, as shown in [37].

In this paper we applied oncotree2vec to mutation trees that contain point mutations. However, our method can be easily expanded to copy number trees and to mutation trees containing any sets of genomic alterations such as mixed trees of joint copy number and point mutations [47], or augmented trees [17]. Learning tree embeddings will facilitate the integration of tumor mutation trees with data from other technologies in downstream analysis (e.g., in deep learning approaches for single-cell multi-omics data integration).

## Acknowledgements

The authors would like to thank Prof. Katharina Jahn for providing the mutation trees, Prof. Koichi Takahashi for providing the clinical data for the AML cohort from [35], and Diane Duroux and Pawewl Czy ż for carefully reading the manuscript.

