## Supplementary information for "oncotree2vec – A method for embedding and clustering of tumor phylogenetic trees"

**Supplementary Table 1: Parameters used in the experiments from Section 3.** The use of the parameters is described in Section 2.2. For the two experiments with not applicable kernel size (NA) we discard the tree vocabulary words corresponding to different neighborhoods, by the design of the experiment. We used WL kernel size 9, equal to the longest path from root to leaves in the cohort, for the first synthetic dataset containing 16 groups of trees, in order to ensure that we match entire tree structures, since the dataset was designed such that the trees from the same group share the same tree structure. The vocabulary categories not listed in the table were discarded.

| Experiment | Max WL kernel size | Vocabulary categories | Augmentation amounts | Embe-dding size ($\delta$) | Num. of training iterations until convergence | Vocabulary size (number of words) | CPU time on a desktop PC |
|---|---|---|---|---|---|---|---|
| Synthetic dataset I | 9 | Neighborhoods<br>Tree structures | 1<br>1 | 128 | 7,000 | 92,784 | 2h31 |
| Synthetic dataset II | NA | Individual nodes | 1 | 64 | 200 | 5,200 | 0m46 |
| Selected pairs of trees, oncotree2vec without neighborhoods | NA | Root-child<br>Direct edges<br>Same path<br>Mutual exclusivity | 1<br>1<br>1<br>1 | 32 | 600 | 60 | 0m51 |
| Selected pairs of trees, oncotree2vec wlk size 1 | NA | Neighborhoods<br>Root-child<br>Direct edges<br>Same path<br>Mutual exclusivity | 1<br>5<br>5<br>5<br>5 | 32 | 600 | 740 | 1m |
| Real trees with different modes of evolution | 3 | Tree structures | 5 | 64 | 500 | 4,906 | 3m34 |
| AML mutation trees | 3 | Neighborhoods<br>Individual nodes<br>Root-child<br>Direct edges<br>Same path<br>Mutual exclusivity | 1<br>5<br>20<br>10<br>10<br>10 | 128 | 1,500 | 33,920 | 1h45 |

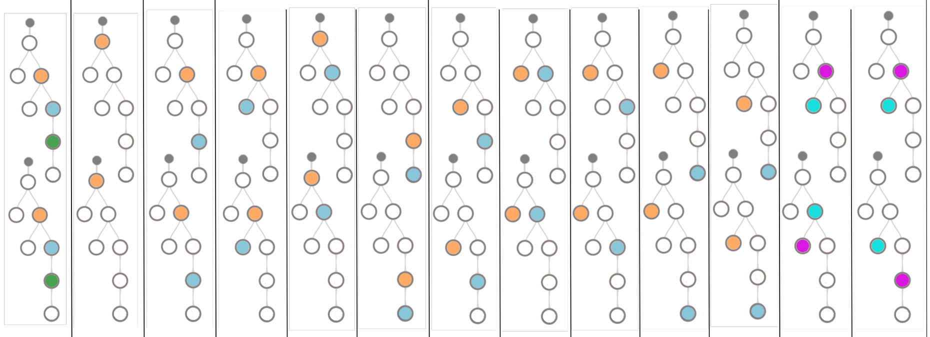

| | pairwise relations | | | | | | | | | | | | | |
|---|---|---|---|---|---|---|---|---|---|---|---|---|---|---|
| | neigh-borhood | root-child | path | direct edges | | | mutual exclusivity | | | | | non-matching relations | | negatives |
| CASet | 0.687 | 0.593 | 0.647 | 0.647 | 0.672 | 0.650 | 0.651 | 0.649 | 0.649 | 0.649 | 0.651 | 0.651 | 0.650 | 1 |
| DISC | 0.776 | 0.912 | 0.850 | 0.822 | 0.856 | 0.789 | 0.799 | 0.829 | 0.803 | 0.780 | 0.787 | 0.842 | 0.820 | 1 |
| MP3 | 0.926 | error | 0.982 | 0.982 | 0.982 | 0.982 | 0.982 | 0.982 | 0.982 | 0.982 | 0.982 | 1 | 1 | 1 |
| Bourque | 0.714 | 0.928 | 0.714 | 0.789 | 0.857 | 0.642 | 0.642 | 0.785 | 0.710 | 0.571 | 0.571 | 0.857 | 0.785 | 1 |
| graph2vec wlk size 1 | 0.31 | 0.39 | 0.42 | 0.37 | 0.41 | 0.32 | 0.43 | 0.39 | 0.39 | 0.39 | 0.36 | 0.44 | 0.45 | 0.51 |
| graph2vec wlk size 2 | 0.34 | 0.45 | 0.5 | 0.41 | 0.39 | 0.39 | 0.42 | 0.46 | 0.46 | 0.45 | 0.47 | 0.41 | 0.45 | 0.52 |
| oncotree2vec without neighborhoods | 0 | 0 | 0 | 0 | 0 | 0 | 0 | 0 | 0 | 0 | 0 | 0.972 | 0.996 | 0.936 |
| oncotree2vec wlk size 1 | 0.19 | 0.23 | 0.27 | 0.24 | 0.21 | 0.19 | 0.3 | 0.28 | 0.3 | 0.26 | 0.27 | 0.52 | 0.48 | 0.57 |

**Supplementary Table 2: Normalized pairwise distance scores between selected pairs of trees, using various distance metrics (synthetic dataset III).** For simplicity, we matched pairs of trees with the same structure and a subset of selected matching node labels. The pairs of matching trees are displayed in the header of the table and the colored nodes correspond to matching nodes in each pair of trees. Each pair contains only one matching pattern, one from each vocabulary category, as described in Fig. 2. The trees match two by two; there is no matching between trees outside the pairs. We add two pairs which do not match because of the reverse order of the mutation events, and the different branching of the events, respectively. We also show the average distance between all the non-matching trees in the tree set (last column "negatives"). The distance scores shown are in the range [0,1] (0 indicates the highest and 1 indicates the lowest similarity). For Bourque distance we use normalized scores and apply it to rooted trees. For MP3, which reports similarity scores between 0 and 1, the values were converted by subtracting 1 from the MP3 score in order to obtain the distance scores. The scores for graph2vec and oncotree2vec are averaged over 10 different runs. The distance metrics considered are CASet and DISC (DiNardo et al. 2020), MP3 (Ciccolella et al. 2021), Bourque (Jahn et al. 2016). For graph2vec (Narayanan et al. 2017) and oncotree2vec we apply the cosine distance to the learned embeddings obtained after an optimal number of iterations, as described in Section 2.2. The colored cells indicate problematic scores, as described in Section 3.1.3.

**Supplementary Table 3: Clusters in the AML mutation tree cohort, sorted by cluster size.** The mean survival time is indicated for clusters of 4 samples or more. The cluster ids correspond to the ones in Fig. 7 and Suppl Fig. 5. The colored rows correspond to groups of clusters (the blue clusters vs the yellow one) with significant difference in the survival curves according to the pairwise log-rank test; the result is confirmed by the values of the median survival time. The NA values indicate that the median survival could not be computed because the survival data did not drop below 50% at the end of the available data (see also the Kaplan Meier curves in Suppl. Fig. 5).

| Number of samples | Shared mutations | Median survival time (number of months) | Cluster id |
|---|---|---|---|
| 18 | DNMT3A primary mutation | 17.9 | 0 |
| 16 | IDH2 primary mutation | 55.3 | 1 |
| 11 | TET2 primary mutation | 11.2 | 2 |
| 9 | NRAS primary mutation | 12.9 | 3 |
| 7 | TP53 primary mutation | 9.3 | 4 |
| 6 | WT1 primary mutation | 12.65 | 5 |
| 6 | SF3B1 primary mutation | 13.9 | 6 |
| 5 | IDH1 primary mutation | 16.7 | 7 |
| 5 | NPM1 primary mutation<br>PTPN11, KRAS clonal exclusivity | 24.9 | 8 |
| 4 | FLT3-ITD primary mutation | 9.1 | 9 |
| 4 | SRSF2 primary mutation | 23.57 | 10 |
| 4 | NPM1 primary mutation | NA | 11 |
| 4 | FLT3 primary mutation<br>FLT3, NRAS clonal exclusivity | NA | 12 |
| 3 | TP53, DNMT3A co-occurrence | - | - |
| 3 | IDH2, DNMT3A, SRSF2 co-occurrence | - | - |
| 2 | PTPN11, NPM1 co-occurrence<br>(identical samples) | - | - |
| 2 | EZH2, ASXL1, RUX1<br>co-occurrence | - | - |
| 2 | IDH2, NPM1, SRSF2<br>co-occurrence | - | - |
| 2 | DNMT3A, KRAS, NPM1, FLT3<br>co-occurrence | - | - |
| 2 | DNMT3A, SF3B1, FLT3, RUNX1<br>co-occurrence | - | - |

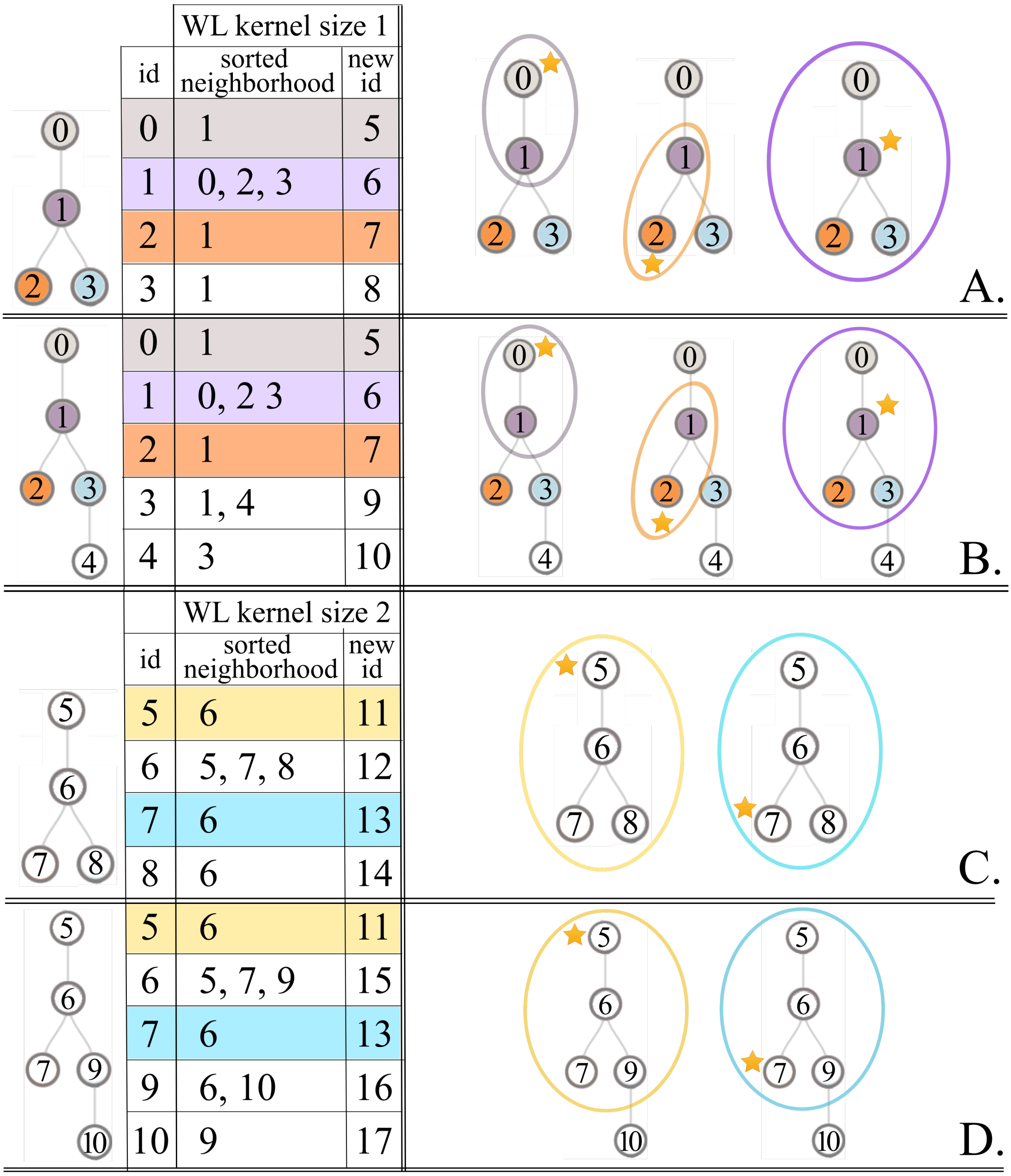

**A.**

| | WL kernel size 1 | |
|---|---|---|
| id | sorted neighborhood | new id |
| 0 | 1 | 5 |
| 1 | 0, 2, 3 | 6 |
| 2 | 1 | 7 |
| 3 | 1 | 8 |

**B.**

| id | sorted neighborhood | new id |
|---|---|---|
| 0 | 1 | 5 |
| 1 | 0, 2 3 | 6 |
| 2 | 1 | 7 |
| 3 | 1, 4 | 9 |
| 4 | 3 | 10 |

**C.**

| | WL kernel size 2 | |
|---|---|---|
| id | sorted neighborhood | new id |
| 5 | 6 | 11 |
| 6 | 5, 7, 8 | 12 |
| 7 | 6 | 13 |
| 8 | 6 | 14 |

**D.**

| id | sorted neighborhood | new id |
|---|---|---|
| 5 | 6 | 11 |
| 6 | 5, 7, 9 | 15 |
| 7 | 6 | 13 |
| 9 | 6, 10 | 16 |
| 10 | 9 | 17 |

**Supplementary Figure 1: Example of neighborhood matching scheme between two trees using the Weisfeiler-Lehman (WL) kernel. (A,B)** For each of the two trees, each node is represented as a sorted list of its first degree neighbors and an id is assigned to each such neighborhood. The new ids encode the neighborhoods of size 1 around each node. There are 3 matching neighborhoods of size 1 between the two trees, around the nodes indicated with stars. Note that a neighborhood of size 1 around the middle node of the first tree covers the entire tree. **(C,D)** A new iteration of the WL subtree kernel is equivalent to encoding neighborhoods of size 2 around every node. The trees are relabeled with the new ids computed in the previous iteration. There are 2 matching neighborhoods of size 1 between the two trees, around the nodes indicated with stars. The colors help matching the tree neighborhood visualization with the corresponding neighborhood labels from the table.

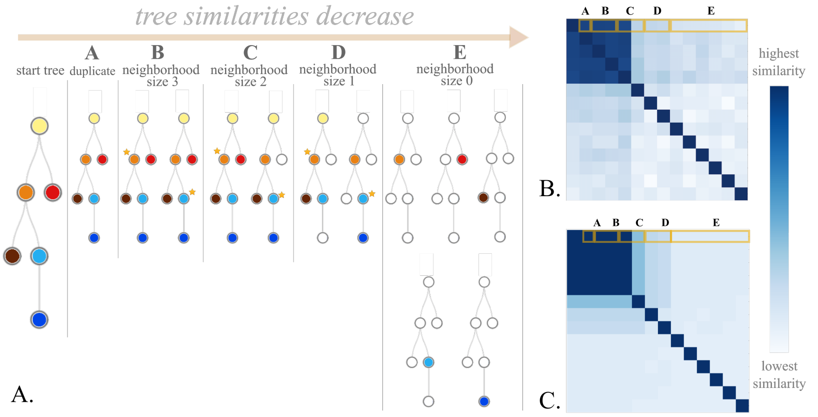


D.

| category | number of vocabulary intersections | rank | similarity w.r.t. start tree | minimum similarity in previous rank | similarity deviation from expected rank |
|---|---|---|---|---|---|
| A | 114 | 1 | 0.9728 | 0.9728 | 0 |
| B | 114 | 1 | 0.9719 | 0.9728 | 0 |
| B | 114 | 1 | 0.9702 | 0.9728 | 0 |
| C | 114 | 1 | 0.9667 | 0.9728 | 0 |
| C | 67 | 2 | 0.7313 | 0.9667 | 0 |
| D | 61 | 3 | 0.7182 | 0.7313 | 0 |
| D | 60 | 4 | 0.7198 | 0.7182 | 1.5e-3 |
| E | 55 | 5 | 0.6895 | 0.7198 | 0 |
| E | 55 | 5 | 0.6837 | 0.7186 | 0 |
| E | 55 | 5 | 0.6823 | 0.7186 | 0 |
| E | 55 | 5 | 0.6802 | 0.7186 | 0 |
| E | 55 | 5 | 0.6690 | 0.7186 | 0 |
| E | 55 | 5 | 0.6514 | 0.7186 | 0 |

**Supplementary Figure 2: Example of simulated mutation trees with decreasing similarities, starting from a given start tree. (A)** Tree group construction: all trees share the same tree structure, but the number of matching nodes is restricted to specific neighborhoods of decreasing sizes around every node with degree greater than 2 (for example, the starred nodes in the figure). Matching nodes are colored the same. In this particular example, a neighborhood of size 3 around a middle node covers the whole tree, leading to an identical match. Nevertheless, a duplicate of the start tree is included in the small cohort from the beginning. The categories A-E denote the different matching neighborhood sizes and the similarities w.r.t. the start tree are expected to decrease as the size of the matching neighborhoods decreases. **(B)** Corresponding tree distance heatmap based on the embeddings learned by oncotree2vec. The heatmap is built using hierarchical clustering with the cosine distance metric on the tree embeddings. **(C)** Heatmap showing the number of shared vocabulary words (encoding neighborhoods of different sizes) for each pair of trees. **(D)** Similarity of each tree w.r.t. the start tree using the learned embeddings (highest similarity is 1). The similarity scores are ordered by the number of vocabulary intersections in decreasing order. The maximum intersection size, 144, corresponds to the identical match to the start tree and is computed as the number of tree nodes times the size of the largest WL kernel applied for the labeled neighborhoods and subtree structures, i.e., $6x(10 + 9)$ – Section 2.1; Suppl. Table 1, synthetic dataset II. Similarity scores corresponding to the same rank are colored the same. Each similarity score should be lower or equal to the minimum score from the previous rank. The deviation of the embedding similarities from the expected order is computed by subtracting, for each tree, the computed similarity w.r.t. the start tree from the minimum similarity score from the previous rank. In this example, there is one embedding deviating from the expected ordering (similarity deviation 0.0015). The average deviation from the expected rank ordering is 0.000782, which is very close to zero, the deviation for perfect ordering.

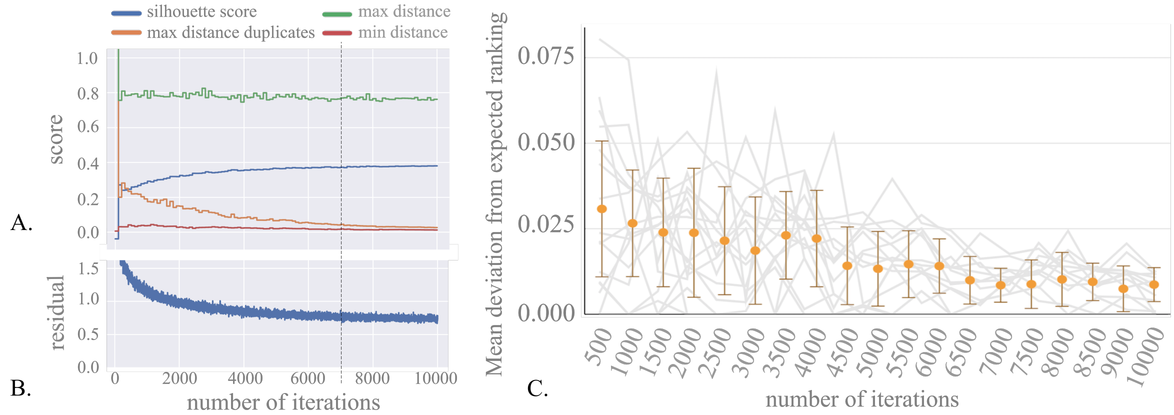


**Supplementary Figure 3: Clustering a cohort of 16 groups of synthetic trees (synthetic dataset I). (A)** Training parameters: minimum and maximum cosine distance between the learned embeddings after every training iteration (see Section 2.2). **(B)** The residual after each training iteration. The dotted line shows where the convergence plot becomes steady, i.e., the training algorithm starts to converge (this is the cutoff used). **(C)** Line chart showing the deviation from the expected rank ordering of the trees in each group. The lines correspond to the scores computed after every 5,000 iterations (10,000 iterations in total). The error bars show the average deviation from the expected rank ordering across all the 16 cohorts and the corresponding standard deviation.

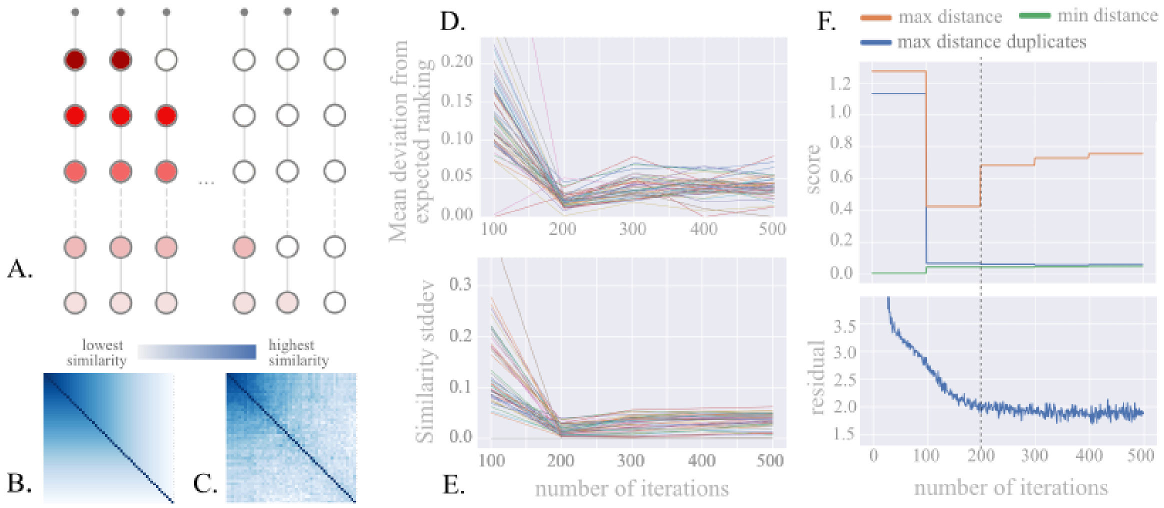


**Supplementary Figure 4: Clustering synthetic trees with known similarity rank ordering (synthetic dataset II). (A)** Cohort of synthetic linear trees with decreasing number of matching nodes. Matching nodes are colored the same. **(B)** Heatmap showing the number of matching nodes for each pair of trees, sorted by the number of matches.This is the expected tree rank ordering. Note that the trees from the antidiagonals parallel to the main antidiagonal share the same number of matching nodes (the pairwise vocabulary intersections have the same number of elements). **(C)** Tree similarity heatmap of the pairwise distances between the simulated trees, using the cosine distance between the learned embeddings. The order of the trees corresponds to the one in panel B. **(D)** Line chart showing the deviation of the embedding similarities from the expected rank ordering with respect to every tree in the cohort, computed every 100 iterations. **(E)** Standard deviation of the pairwise tree embedding distances between the learned tree embeddings corresponding to pairs of trees with the same number of matching nodes, computed every 100 iterations. Each line corresponds to the pairs of trees that share a number of matching nodes ranging from 0 to 50, which appear on the same antidiagonal in panel B. **(F)** Training parameters: minimum and maximum cosine distance between the learned embeddings (see Methods 2.2) and the residual after each training iteration. The dotted line shows where the convergence plot becomes steady, i.e., the training algorithm starts to converge (this is the cutoff used).

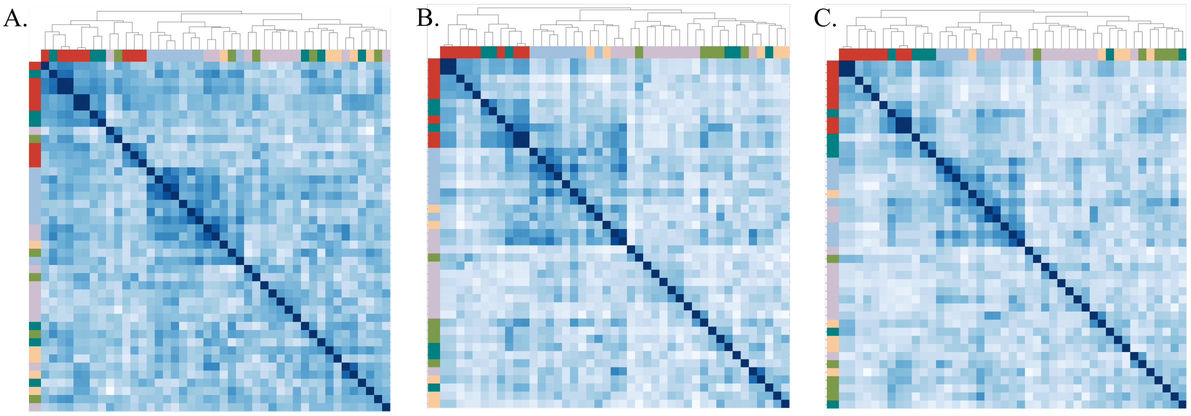


**Supplementary Figure 5: Result of clustering tree structures from six cancer types** from Noble et al. 2022 using embedding sizes of 32 (A), 64 (B) and 128 (C).

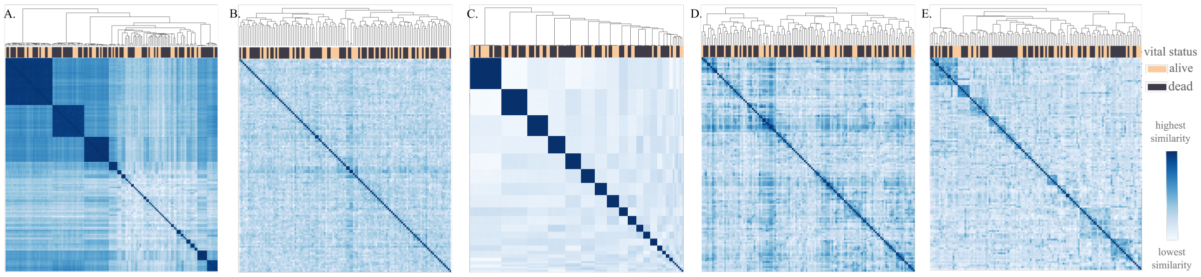


**Supplementary Figure 6:** Hierarchically-clustered heatmap of tree similarities for the AML mutation trees from Morita et al. 2020 using different vocabulary augmentation amounts. **(A)** vocabulary contains only on the tree structures (the labels are discarded); **(B)** vocabulary contains neighborhoods of all sizes up to size 3 – this would be the result obtained using graph2vec; **(C)** vocabulary contains only root child relations; **(D)** vocabulary contains individual nodes, pairwise relations and mutually exclusive pairs; **(E)** vocabulary is based on all the subtree structure categories described in Fig. 2, with the category of pairwise-relations overrepresented (augmented).

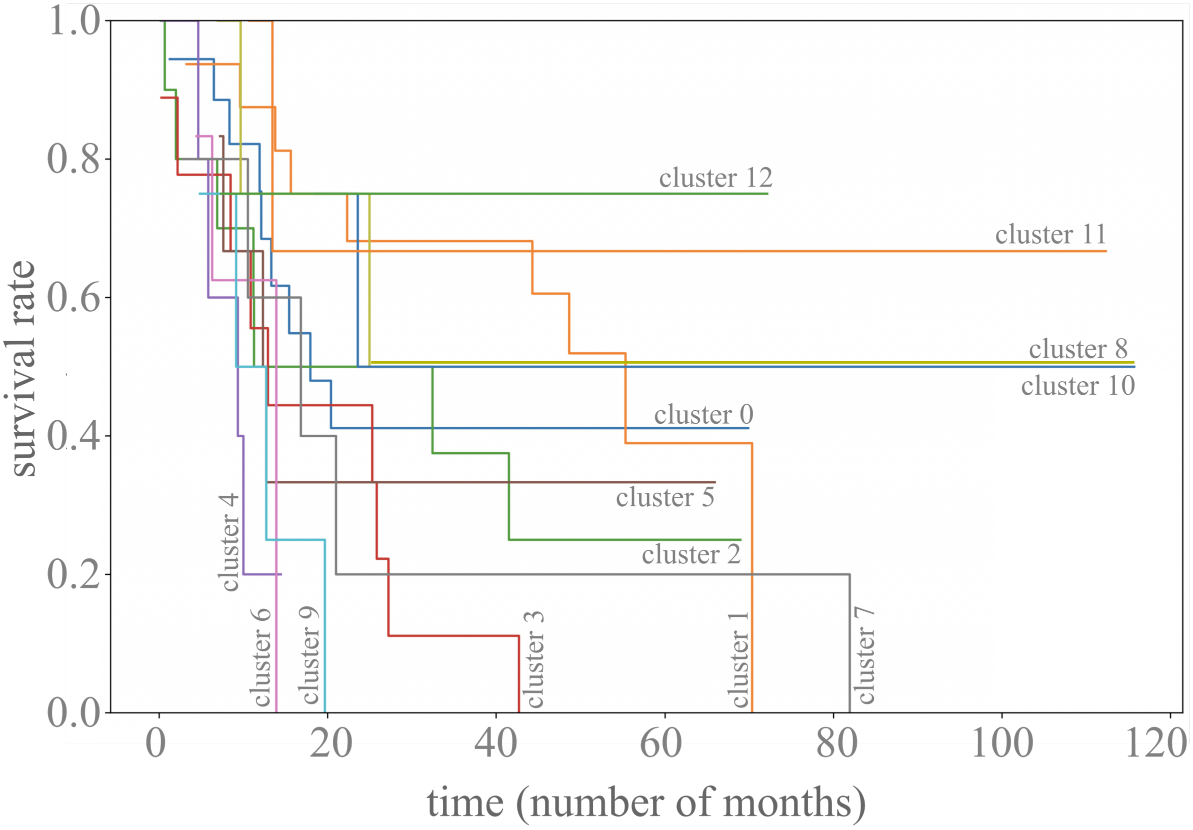


**Supplementary Figure 7: Kaplan Meier curve estimation** of the survival for 13 clusters found in the AML mutation tree cohort (Section 3.3).
